# Structural basis for selective targeting of Rac subfamily GTPases by a bacterial effector protein

**DOI:** 10.1101/2020.06.29.167221

**Authors:** Nikolaus Balthasar Dietz, Markus Huber, Isabel Sorg, Arnaud Goepfert, Alexander Harms, Tilman Schirmer, Christoph Dehio

## Abstract

Ras-homology (Rho) family GTPases are conserved molecular switches controlling fundamental cellular activities in eukaryotic cells. As such, they are targeted by numerous bacterial toxins and effector proteins, which have been intensively investigated regarding their biochemical activities and discrete target spectra; however, molecular mechanisms of target selectivity have remained elusive. Here, we report a bacterial effector protein that targets all four Rac subfamily members of Rho family GTPases, but none of the closely related Cdc42 or RhoA subfamilies. This exquisite target selectivity of the FIC domain AMP-transferase Bep1 from *Bartonella rochalimae* is based on electrostatic interactions with a subfamily-specific pair of residues in the nucleotide-binding motif and the Rho insert helix. Residue substitutions at the identified positions in Cdc42 facilitate modification by Bep1, while corresponding Cdc42-like substitutions in Rac1 greatly diminish modification. Our study establishes a structural paradigm for target selectivity towards Rac subfamily GTPases and provides a highly selective tool for their functional analysis.

## Introduction

Small GTPases of the Ras-protein superfamily are molecular switches that control fundamental cellular functions in eukaryotes by cycling between GTP-bound “on” and GDP-bound “off” conformational states of their switch regions 1 (Sw1) and 2 (Sw2) (Didsbury et al., 1989; Wennerberg et al., 2005). Members of the Ras-homology (Rho) protein family function as signaling hubs and regulate cytoskeletal rearrangements, cell motility, and the production of reactive oxygen species (Heasman and Ridley, 2008; Jaffe and Hall, 2005). Rho family GTPases are defined by the presence of the highly variable, 13 residues long, α-helical Rho insert close to the C-terminus that has been implicated in wiring Rho family GTPases to their specific biological functions (Bokoch and Diebold, 2002; Karnoub et al., 2004). Additionally, the otherwise invariable TKxD Ras-nucleotide-binding motif is altered to TQxD in a subset of Rho family GTPases.

Due to their central role in eukaryotic cell signalling Rho family GTPases are targeted by a plethora of bacterial virulence factors, including secreted bacterial toxins that autonomously enter host cells and effector proteins that are directly translocated from bacteria into host cells via dedicated secretion systems (Aktories, 2011, 2015). By means of these virulence factors, pathogens established ways to stimulate, attenuate or destroy the intrinsic GTPase activity of Rho family GTPases, either directly through covalent modification of residues in the Sw1 or Sw2 regions (Aktories, 2015), or indirectly by mimicking guanine nucleotide exchange factor (GEF) or GTPase-activating protein (GAP) function. However, the structural basis for selective targeting of Rho family GTPase subfamilies has remained unknown (Aktories, 2011).

The bacterial genus Bartonella comprises a rapidly expanding number of virtually omnipresent pathogens adapted to mammals, many of which have been recognized to cause disease in humans (Wagner and Dehio, 2019). The stealth infection strategy of *Bartonella* spp. (Harms and Dehio, 2012) relies to a large extend on translocation of multiple Bartonella effector proteins (Beps) via a dedicated type IV secretion system. Strikingly, the majority of the currently known several dozens of Beps contains enzymatic FIC domains (Harms et al., 2017; Wagner and Dehio, 2019), indicating that *Bartonella* spp. successfully utilize this effector type in their lifestyle. In order to gain more insights into the function of FIC domain-containing Beps we have here investigated Bep1 of *Bartonella rochalimae* originally described by (Harms et al., 2017).

FIC (filamentation induced by cyclic AMP) domain-containing effector proteins represent a family of ubiquitous proteins with a conserved molecular mechanism for post-translational modification of target proteins. FIC domains are comprised of six helices with a common HxFx(D/E)GNGRxxR motif between the central helices 4 and 5 (Harms et al., 2016). Some of the FIC domain-containing effector proteins have been recognized to modify Rho family GTPases by catalyzing transfer of the AMP-moiety from the ATP substrate to specific target hydroxyl side-chains (reviewed in Harms et al., 2016; Hedberg and Itzen, 2015). Prototypical examples are the effector proteins IbpA from *Histophilus somnii* and VopS from *Vibrio parahaemolyticus*, which both target a wide range of Rho family GTPases and AMPylate (adenylylate) a conserved tyrosine or threonine residue of Sw1, respectively (Mattoo et al., 2011; Worby et al., 2009; Yarbrough et al., 2009). Both modifications result in abrogation of downstream signaling, causing collapse of the cytoskeleton of the host cell and subsequent cell death (Roy and Cherfils, 2015). Here, we show that the FIC domain of Bartonella effector protein 1 of *Bartonella rochalimae* (Bep1) AMPylates the same Sw1 tyrosine residue as IbpA, while the target spectrum is strictly limited to the Rac subfamily of Rho GTPases. Employing a combination of structural analysis, modelling, biochemistry, and mutational analysis, we identify the structural determinants of this remarkable target selectivity. Our findings highlight the potential of Bep1 as a novel tool for dissecting Rho family GTPase activities and provide a rationale for the re-design of its target selectivity.

## Results

### Bep1 selectively AMPylates Rac subfamily GTPases

Bep1 is composed of a canonical FIC domain followed by an oligosaccharide binding (OB) fold and a C-terminal BID domain (Harms et al., 2017). Latter domain is implicated in recognition and translocation by the type 4 secretion system VirB/VirD4 of Bartonella (Schulein, Guye et al., 2005; Wagner et al., 2019).

In search for Bep1 targets we performed AMPylation assays by incubating lysates of *E. coli* expressing Bep1 with eukaryotic cell lysates and α-^32^P-labelled ATP and observed a radioactive band migrating with an apparent molecular weight of 20 kDa (Fig. S1A), consistent with modification of Rho family GTPases as previously described for IbpA and VopS (Worby et al., 2009; Yarbrough et al., 2009). To investigate further, we explored the target spectrum of Bep1 and compared it to those of the FIC domains of IbpA (IbpA_FIC2_) or VopS (VopS_FIC_) by selecting 19 members of the Ras superfamily (Fig. 1A) with an emphasis on members of the Rho family. While AMPylation activity of all three enzymes was strictly confined to Rho family GTPases, their target selectivity spectra differed markedly: While Bep1 modified exclusively members of the Rac subfamily (i.e., Rac1/2/3, and RhoG), the target spectrum of IbpA_FIC2_ comprised all Rho GTPases with the exception of RhoH/U/V and the Rnd subfamily, and VopS_FIC_ was found to be fully indiscriminative (Fig. 1A, summarized in D).

**Figure 1.**
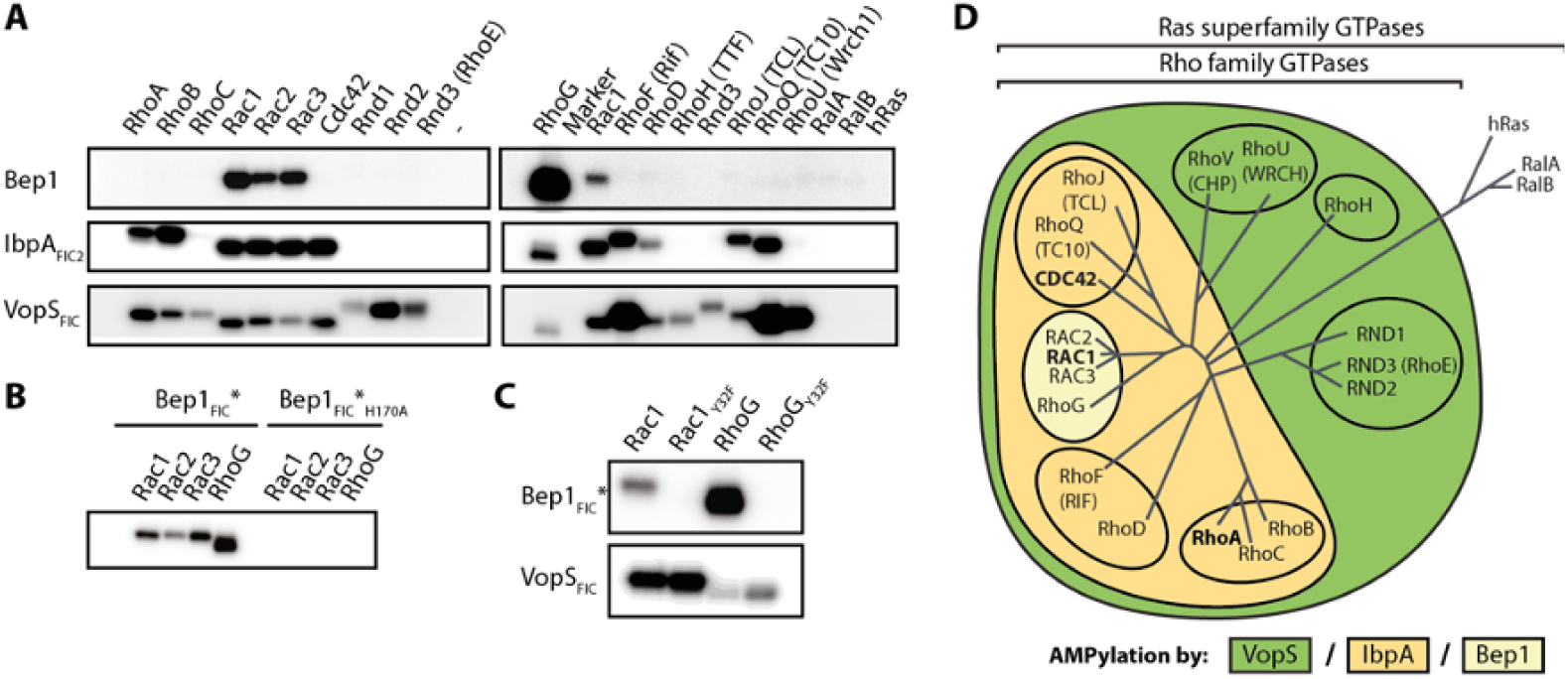
Bep1 selectively targets Rac subfamily GTPases. (**A**) ^32^P-autoradiograms of *in vitro* AMPylation reactions using the indicated purified and GDP-loaded Rho family GTPases display exquisite selectivity of full-length Bep1 for Rac subfamily GTPases in contrast to the broader target spectrum of IbpA_FIC2_ and VopS_FIC_. (**B**) The FIC domain of Bep1 in complex with the regulatory protein BiaA (Bep1_FIC_*) is sufficient for the recognition of Rac subfamily GTPases and the catalytic H170 is required for AMPylation. (**C**) Bep1_FIC_* AMPylates residue Y32 of Rac1 and RhoG, since the respective Y32F mutants are not modified. AMPylation by the T35-specific VopS_FIC_ indicates structural integrity of the analyzed GTPases and their Y32F mutants. (**D**) Venn diagram showing AMPylation target selectivity of tested FIC domains, overlaid to the phylogenetic relation of Rho-family GTPases (Heasman and Ridley, 2008).

Next, we designed a minimal Bep1_FIC_ construct (residues 13 – 229) that proved sufficient for selective target modification. Bep1 belongs to the class I of FIC proteins that are regulated by a small regulatory protein, here BiaA, that inhibits FIC activity by inserting a glutamate residue (E33) into the ATP binding pocket (Engel, Goepfert et al., 2012). In order to improve expression level and stability, we co-expressed Bep1_FIC_ with an inhibition relieved mutant (E33G) of BiaA, yielding the stabilized minimal AMPylation-competent Bep1_FIC_/BiaA_E33G_ complex, in short Bep1_FIC_*

Bep1_FIC_* efficiently AMPylates its targets and the activity depends on the presence of the catalytic histidine (H170) of the signature motif (Fig. 1B), consistent with the canonical AMPylation mechanism (Engel, Goepfert et al., 2012). Bep1_FIC_*, in contrast to VopS_FIC_, does not AMPylate Rac1_Y32F_ (Fig. 1C), indicating that Bep1_FIC_* modifies Y32 of the Rac1 Sw1 as confirmed by mass spectrometry (Fig. S1C). Thus, Bep1_FIC_* catalyzes the equivalent modification as IbpA_FIC2_ (Worby et al., 2009; Xiao et al., 2010), whereas VopS modifies T35 (Yarbrough et al., 2009). In contrast to the GDP-form, GTP-loaded GTPases may not be amenable to FIC-mediated modification of Y32 since this residue is known to be involved in GTP binding via interaction with the γ-phosphate group (Lapouge et al., 2000) (Fig. S2D). Indeed, exchanging GDP against GTP efficiently protected the GTP hydrolysis deficient mutant Rac1_Q61L_ from modification and the same effect was observed, when replacing GDP bound to wild-type Rac1 with non-hydrolysable GTPγS (Fig. S2C). Thus, we conclude that GDP-loaded GTPases are the physiological targets of Bep1-mediated AMPylation.

### The crystal structure of Bep1_FIC_* reveals an extended target recognition flap

To reveal the structural basis of target selectivity, we solved the crystal structure of Bep1_FIC_* to 1.6Å resolution. The structure (Fig. 2) closely resembles those of other FIC domains with AMPylation activity such as VbhT (Engel, Goepfert et al., 2012), IbpA (Xiao et al., 2010), and VopS (Luong et al., 2010), featuring the active site defined by the conserved signature motif encompassing the α4-α5 loop and the N-terminal part of α5. Comparison with the apo crystal structure of the close Bep1 homolog from *B. clarridgeiae* (PDB ID 4nps) shows that the presence of the small regulatory protein BiaA in Bep1_FIC_* does not affect the structure of the FIC domain (Fig. S2B).

**Figure 2.**
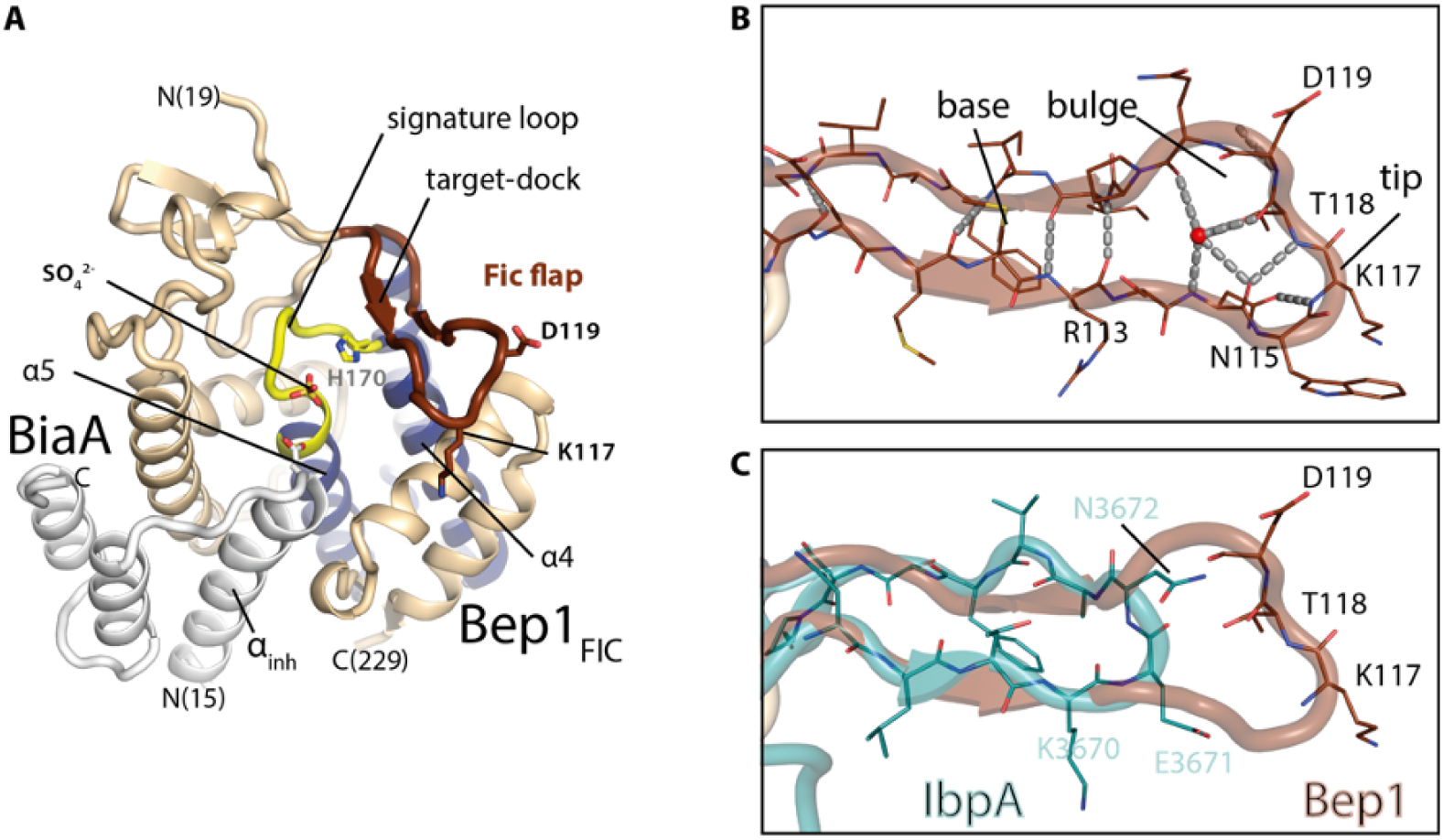
Crystal structure of Bep1_FIC_* reveals extended flap. (**A**) Cartoon representation of the crystal structure of the Bep1_FIC_:BiaA complex (Bep1_FIC_*) determined in this work. The regulatory protein BiaA is shown in light grey. The FIC domain fold is shown in light brown, with the central FIC helices (α4-α5) in blue. The FIC signature loop with the catalytic H170 is shown in yellow, the FIC flap covering the active site in dark brown. (**B**) Detailed view of the Bep1 flap region (PDB ID 5eu0, this study). Structural flap elements are stabilized by an H-bonding network involving main-chain and side-chain groups. H-bonds are shown by grey dashed lines. The base of the flap forms a two-stranded β-sheet, with the N-terminal part constituting the target dock. The tip of the flap forms an i > i+3 turn between N115 and T118, which is further stabilized by the sidechain of N115. The tip is followed by a bulge and a conserved proline residue and stabilized by interactions of the backbone with a central water (in red). This arrangement suggests, that the well-defined structure of the flap orients sidechains K117 and D119 for target interaction. (**C**) Overlay of flaps from Bep1_FIC_ (brown) and IbpA_FIC2_ (turquoise). Residues at the tip of both flaps are indicated. Compared to Bep1, the flap of IbpA is 6 residues shorter amounting to 8Å (see Fig. S2A).

The active site is partly covered by a β-hairpin ‘flap’ (Fig. 2A) that serves to register the segment carrying the modifiable side-chain (here Sw1) to the active site via β-sheet augmentation, as has been inferred from bound peptides (Goepfert et al., 2013; Yarbrough et al., 2009), observed directly in the IbpA_FIC2_:Cdc42 complex (Xiao et al., 2010), and discussed elsewhere (Roy and Cherfils, 2015). Strikingly, the flap of Bep1 and its orthologs in other Bartonella species (Fig. S2A) is considerably longer than in other FIC structures (e.g., of IbpA_FIC2_) and features a well-defined bulge at its tip (Fig. 2B, C).

### Bep1_FIC_:target model suggests that charged residues of the flap determine target selectivity

The complex structure of a FIC enzyme with a small GTPase target and the mechanism of FIC catalyzed AMPylation reaction has been elucidated for IbpA_FIC2_ in complex with GDP-loaded Cdc42 (Xiao et al., 2010) (Fig. 3B). The detailed view in Fig. 3D shows that the Sw1 segment of Cdc42 exhibits an extended conformation and forms antiparallel, largely sequence independent, β-sheet interactions with the flap of the FIC enzyme, thereby aligning the modifiable Y32 with the active site. Considering the close structural homology of the catalytic core of Bep1_FIC_ with IbpA_FIC2_ (rmsd = 1.0Å for 32 Cα atoms in the active site helices) and of Rac subfamily GTPases with Cdc42 (0.44Å /175), we reasoned that computational assembly of a Bep1_FIC_:Rac complex could provide a structural basis for an understanding of Bep1 target selectivity.

**Figure 3.**
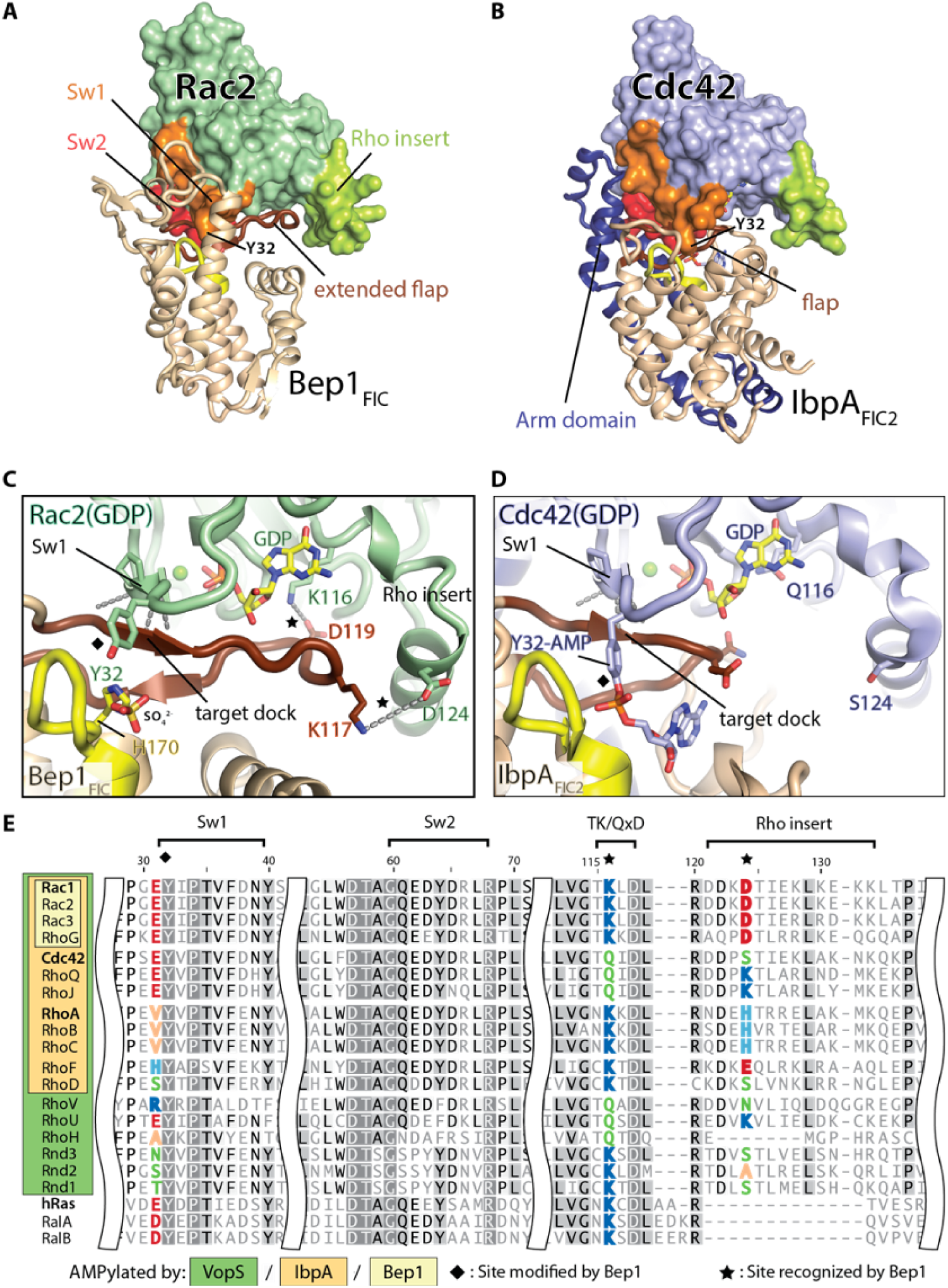
Bep1_FIC_:Rac2 complex model suggests charged interactions between FIC flap and targets. (**A**) Bep1_FIC_:Rac2 complex model and (**B**) IbpA_FIC2_:Cdc42 crystal structure (PDB 4itr). The FIC fold is shown in light brown. The FIC signature loop with the catalytic H170 is shown in yellow, the FIC flap covering the active site in brown. GTPases are shown as surface representation with indicated structural elements distinguished by color: Switch 1 (Sw1) in orange, Switch 2 (Sw2) in red and Rho-insert in green. The extension of the Bep1_FIC_ flap is accommodated in a groove formed by the T(K/Q)xD motif and the Rho insert (**B**), whereas the arm domain of IbpA (in blue) contacts the effector binding regions, Sw1 and Sw2, of the GTPase. (**C, D**) Comparison of intermolecular interactions in (**C**) the Bep1_FIC_:Rac2 model and (**D**) the IbpA_FIC2_:Cdc42 complex. H-bonding and electrostatic interactions are indicated by dashed lines in grey. The tip of the Bep1_FIC_ flap is accommodated in a groove, with K117 and D119 in favorable position to interact with D124 and K116 of Rac2, respectively. (**D**) In the IbpA_FIC2_:Cdc42 complex the Rho insert region is not involved in such interaction. (E) Structure-guided sequence alignment of the GTPases of the Rho, Ras, and RalA/B families. The K116/D124 configuration (marked with a star) is unique to Rac1/2/3 and RhoG (light yellow). Residue numbers refer to Rac1, names of representative members of Rho subfamilies are indicated in bold.

Fig. 3A shows the assembled Bep1_FIC_:Rac2 complex that was obtained by individual superposition of (1) the Bep1_FIC_ active site helices and the flap with the corresponding elements in IbpA_FIC2_ and (2) of the Sw1 loop of Rac2 with that of Cdc42. Thereby, we assumed implicitly that the interaction between these central segments should be very similar, since both FIC enzymes utilize a homologous set of active residues to catalyze AMP-transfer to a homologous residue (Y32) on Sw1.

The local structural alignment resulted in a virtually identical relative arrangement of the FIC core to the GTPase as in the template structure (compare Figs. 3A and B) and caused no steric clashes. Conspicuously, the extended Bep1_FIC_ flap is accommodated in a groove formed by Sw1, the nucleotide binding T(K/Q)xD motif, and the following Rho-insert helix (Fig. 3C; Fig. S2E).

The manually created complex model was used as input for an adapted Rosetta-modelling protocol to allow for sampling of backbone and side-chain torsion angles in the interface of the complex, as described in the method section (Barlow et al., 2018; Kapp et al., 2012). Consistent with the low affinity of the complex *in vitro* (see below), the models confirm the relatively small interface area of approximately 800Å^2^. Common to all top scoring models we find that the modifiable residue Y32 is pointing towards the active site of Bep1, where it is held in place by a mainchain-mediated interaction between the base of the flap and the Sw1 loop of the GTPase (Fig. S3A), indicating that the configuration of active site residues and the modifyable tyrosine side-chain is, indeed, most likely the same as in the template complex.

However, in the IbpA_FIC2_:Cdc42 complex, the aforementioned GTPase groove on the nucleotide binding face is not utilized for the contact (Fig. 3D). Instead, the so called arm domain of IbpA_FIC2_ (Fig. 3B) constitutes a major part of the interface and contacts the highly conserved Sw2 loop of Cdc42. This rationalizes the broad target spectrum of arm domain-containing FIC AMP-transferases like IbpA and VopS (Harms et al., 2016; Luong et al., 2010). In turn, residues of the groove predicted to get recognized exclusively by Bep1_FIC_ are likely to be important for the limited target range of Bep1. Conspicuously, the top scoring models revealed two potential salt-bridges between the Bep1 flap and the Rac2 groove, namely D119(Bep1) - K116(Rac2) and K117(Bep1) - D124(Rac2) (Fig. 3C, S3A). Since the combination of K116 and D124 is exclusively found in the Rac sub-family as revealed by sequence alignment of Rho family GTPases (Fig. 3E), we reasoned that these residues may contribute significantly to the specific recognition of Rac GTPases by Bep1 (Fig. 1A).

### Two salt-bridges between flap and target are crucial for selective interaction of Bep1_FIC_ with Rac subfamily GTPases

The relevance of the two identified salt-bridges in the Bep1_FIC_*:Rac2 complex (Fig. 3C) for affinity and selectivity was tested by single- and double replacements of the constituting residues 116 and 124 in a Bep1 target and a non-target GTPase. For Rac1, we tested if substitutions at these residues with corresponding amino acids of Cdc42 - a non-target of Bep1 with the highest conservation in regions flanking the proposed interaction sites (Fig. 3E) - influence target recognition (loss-of-function approach, see interaction schemes in Fig. 4A). In addition, we tested whether Cdc42 can be converted to a Bep1 target by reciprocal substitution(s) of these sites with the corresponding Rac1 residues (gain-of-function approach, Fig. 4B).

**Figure 4.**
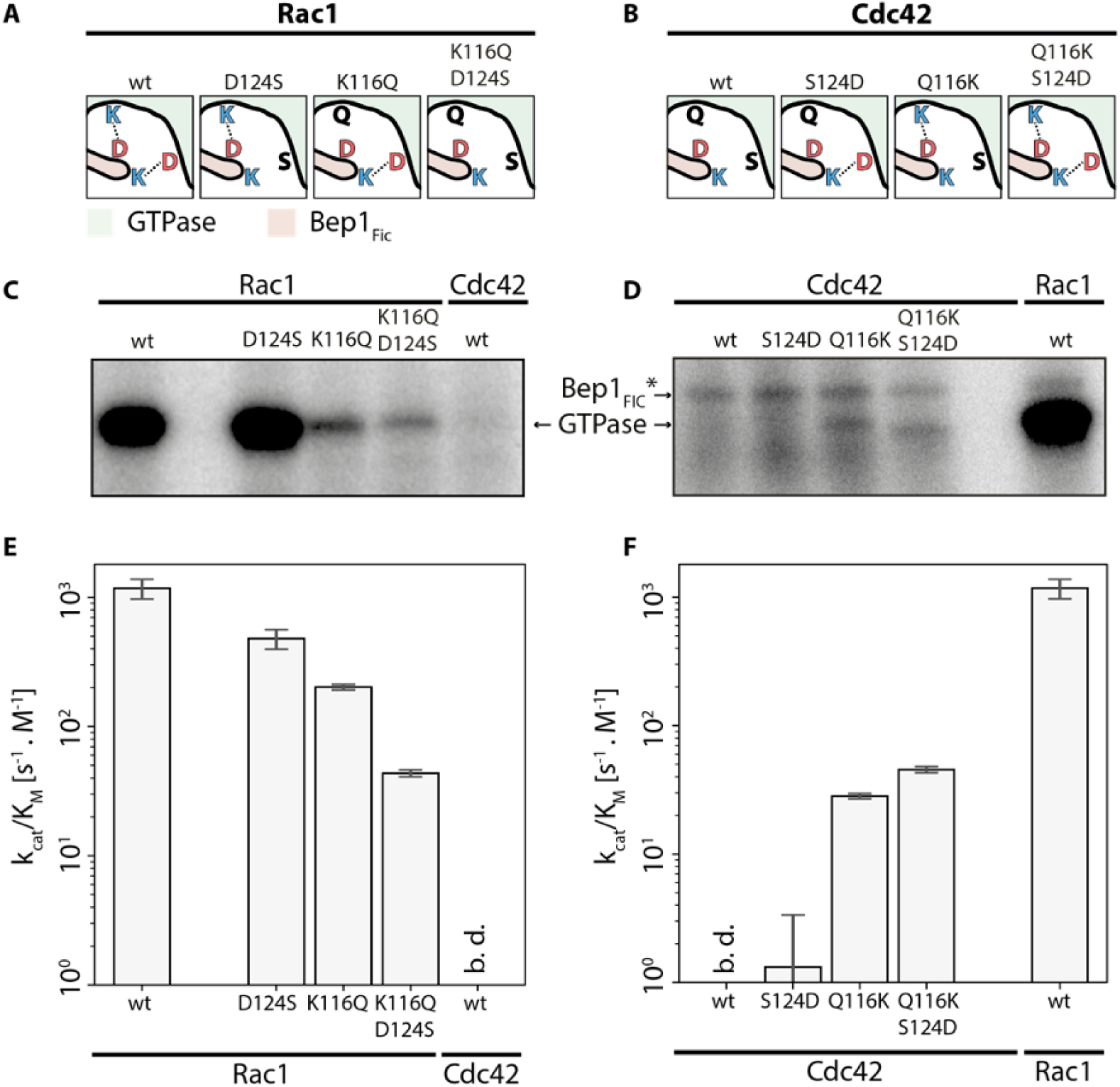
Two salt-bridges are crucial for Rac subfamily-selective AMPylation. (**A**) Schematic view of the two intramolecular Bep1_FIC_:Rac1 salt-bridges (left) and their partial disruption upon site-directed Rac1 mutagenesis, yielding Rac1 loss-of-function mutants (right). (**B**) Absence of ionic interactions in the predicted Bep1_FIC_:Cdc42 interface (left) and partial establishment of salt-bridges in Cdc42 gain-of-function mutants (right). (**C, D**) AMPylation of the variants given in panels (**A, B**) as measured by autoradiography. Note that, due to the employed higher Bep1_FIC_* concentration (see Material and Methods), the experiments in panel D also revealed auto-AMPylation of Bep1_FIC_*. (**E, F**) Enzymatic efficiency constants, k_cat_/K_M_, for Bep1_FIC_* catalyzed AMPylation of the GTPase variants shown in (**A, B**) as derived from the oIEC measurements shown in Figure S4. b.d. – below detection limit.

First, we applied, as for Fig. 1A, the autoradiography end-point assay with [α-^32^P]-ATP as substrate. Compared to wild-type Rac1, mutant D124S showed no significant difference in the amount of AMPylated target, whereas AMPylation of mutant K116Q and, even more, of the double mutant was found drastically reduced (Fig. 4C, S4A). Conversely, in the gain-of-function approach, Cdc42 mutant S124D did not convert the GTPase to a Bep1 target, while mutant Q116K mutant and the double mutant showed low, but significant AMPylation (Fig. 4D, S4B). In a fairly undiscriminating way, IbpA_FIC2_ modified all investigated GTPase variants (Fig. S4C,D) indicating their proper folding. Together, the semi-quantitative radioactive end-point assay demonstrated a major role of K116 in target recognition by Bep1_FIC_*, while a contribution of D124 could not be demonstrated.

To overcome the limitations of the radioactive end-point assay and to characterize target AMPylation quantitatively, we developed an online ion exchange chromatography (oIEC) assay (see Methods) which allows efficient acquisition of enzymatic progress curves to determine initial velocities, v_init_ (see for instance inset to Fig. S4F). For AMPylation of Rac1 by Bep1_FIC_*, titrations experiments yielded 0.52 mM and 1.4 mM for the substrates ATP and Rac1, respectively, and a k_cat_ of 1.9 s^-1^. The comparison with published values on other Fic AMP transferases (Tab. S1) shows that the K_M_ values are comparable to IbpA, but that k_cat_ is smaller by about two orders of magnitude.

Considering the physiological conditions in the cell with an ATP concentration above K_M_, Bep1 can be expected to be saturated with ATP and only partially loaded with the target ([*target*] << *K*_*M,target*_). In such a regime, the AMPylation rate will be given by

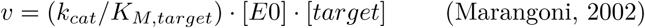

i.e., will depend solely on the second order rate constant k_cat_/K_M,target_ (efficiency constant), which is, thus, the relevant parameter for enzyme comparison. Next, we determined the efficiency constants for all GTPase variants. In the loss-of-function series, the single mutants reduced the efficiency constant by 2- and 6-fold, the double mutant by about 30-fold (Fig. 4E, Table S1). In the gain-of-function series, wild-type Cdc42 showed no and mutant S124D only marginal modification, while mutant Q116K showed a very significant (about 30-fold larger than that of S124D) effect. Again, as in the previous series, the double mutant showed the largest effect that was greater than the sum of the two single mutant contributions (Fig. 4F, Table S1). Summarizing, the quantitative oIEC assay confirmed the prominent dependence of Bep1_FIC_* catalyzed target modification on the type of residue in target position 116 that had already been revealed by the radioactive endpoint assay and predicted by modeling (Fig. S3A), but also demonstrated a significant influence of the residue in position 124. Importantly, the activity data indicate a significant synergistic effect of the two salt bridges on reaction efficiency, which leads to a more than additive decrease or increase of the efficiency constant in the double mutants. Thus, for a substantial change of Bep1 AMPylation efficiency both salt-bridges appear to be crucial.

## Discussion

Although many protein interaction surfaces of Rho family GTPases have been described (Dvorsky and Ahmadian, 2004; Vetter and Wittinghofer, 2001) the basis of discrimination between these structurally conserved but functionally diverse GTPases remained elusive, especially with regard to the role of the highly divergent Rho-insert element. Our structure-function analysis establishes a molecular paradigm of target selectivity for Rac subfamily GTPases that is encoded by intermolecular interaction with the Rho-family specific structural elements. We anticipate that similar mechanisms may be also used by endogenous Rho family GTPase interacting proteins in the physiological context.

We speculate that in the infection process of *Bartonella* spp., the selective inactivation of Rac subfamily GTPases plays a critical role for specifically evading the innate immune response, without causing the collateral damage and activation of the immune system associated with broad-spectrum Rho-GTPase toxins like VopS or IbpA.

Patients with impaired signaling of Rac-subfamily GTPases cannot clear bacterial infections due to diminished ability for ROS production in immune cells, as seen in patients suffering from chronic granulomatosis disease (CGD) or case studies from patients with dysfunctional Rac2 genes resulting in neutrophil immunodeficiency syndrome (NEUID) (Ambruso et al., 2000; Kurkchubasche et al., 2001). At the same time, Rac subfamily-selective AMPylation does not trigger a response of the innate immune system via activation of the pyrin inflammasome, which has been shown to accompany inactivation of RhoA by covalent modification in the Sw1 region (Xu et al., 2014), potentially providing a substantial benefit for *Bartonella* spp. in order to establish a largely asymptomatic chronic infection in their host. Along these lines, we speculate that selective targeting of GDP complexed Rac subfamily GTPases provides the additional benefit that protein levels of GDP-bound Rac are not down-regulated via proteasomal degradation (Lynch et al., 2006), allowing to build a stable pool of inactive Rac subfamily GTPases that would subdue Rac-mediated immune responses effectively.

Beyond providing a molecular paradigm for target selectivity among Rho family GTPases, the narrow target spectrum of Bep1 for Rac subfamily GTPases also provides a unique tool for dissecting their specific functions in cellular processes, such as cytoskeletal rearrangements related to the Rac1-dependent formation of membrane ruffles, the Rac2/RhoG-dependent production of reactive oxygen in immune cells, or the role of Rac1 in carcinogenesis.

Considering the simple topology and small size of the FIC domain, we find a surprisingly modular division of functions. While the conserved catalytic core allows efficient AMPylation of a target hydroxyl residue located in an extended loop that registers to the active site via β-strand augmentation, target affinity and thereby selectivity is encoded separately in a short loop insertion. The modular nature and amenable size of this structural framework appears well suited for the rational design of synthetic Rho subfamily selective FIC domain AMP-transferases with novel physiological activities and beyond.

## Acknowledgements

We thank the laboratories of Klaus Aktories, Kim Orth, and Seema Mattoo for providing expression constructs of Rho-family GTPases, VopS, and IbpA, respectively. We thank the beamline staff at the Swiss Light Source in Villigen for expert help in data acquisition. Rosetta modelling calculations were performed at sciCORE (http://scicore.unibas.ch/) scientific computing core facility at University of Basel. This work was supported by grants 310030A_173119 from the Swiss National Science Foundation (SNSF, www.snf.ch) and advanced grant 340330 to (FicModFun) from the European Research Council (ERC), both to C.D.

## Author contributions

N.D., M.H., T.S. and C.D. designed research and wrote the paper with contributions from all authors. A.H. initiated and designed the project, cloned expression constructs and performed AMPylation assays. I.S. designed research, cloned and purified expression constructs of GTPases and FIC domain proteins and performed AMPylation assays. A.G co-expressed, crystallized, and solved the structure of Bep1_FIC_:BiaA. N.D. built, refined, analysed and modelled the structure of Bep1 and the Bep1 target complex. N.D. and M.H. cloned, expressed purified Bep1 and mutant GTPases for AMPylation assays. M.H. performed autoradiography assays and time-resolved ion exchange chromatography with GTPase variants and processed the data.

## Data and Material Availability

Coordinates and structure factors for the Bep1_FIC_:BiaA complex structure (Bep1_FIC_*) have been deposited in the Protein Data Bank under accession number 5eu0. Correspondence and requests for materials should be addressed to C.D.

## Star*Methods

### Contact for reagent and resource sharing

Requests for further information and for resources and reagents should be directed to and will be fulfilled by the Lead Contact Christoph Dehio.

### Method details

#### Protein expression and purification

The FIC domain of Bep1 was cloned, expressed and purified in complex with the inhibition-relieved regulatory protein BiaA_E33G_ as described for the crystallization construct and is subsequently referred to as Bep1_FIC_*. For the generation of cleared bacterial lysate, the bacterial pellet was resuspended in reaction buffer (50 mM Tris-HCl pH 8.0, 150 mM NaCl, 5 mM MgCl_2_) supplemented with protease inhibitor cocktail (complete EDTA-free mini, Roche) and lysed by sonication. After clearing the lysates by centrifugation (120’000 g for 30 min. at 4°C), the supernatant was directly used in the assays or stored at −20°C. Protein expression and purification of GST- or HIS-tagged GTPases and GST-tagged FIC domains of VopS and IbpA followed standard GST- or HIS-fusion-tag protocols. In short: *E. coli* BL21 or BL21 AI (Invitrogen) were transformed with expression plasmids and used for protein expression. Bacteria were grown in LB medium supplemented with appropriate antibiotic on a shaker until A600 = 0.6 to 0.8 at 30°C. Protein expression was induced by addition of 0.2 mM isopropyl-β-D-thiogalactopyranoside (IPTG) (AppliChem GmbH, Darmstadt, Germany) or 0.1% w/v arabinose (Sigma-Aldrich, Germany) for 4-5 h at 22°C.

Bacteria were harvested by centrifugation at 6’000 g for 6 min. at 4°C, resuspended in lysis buffer (20 mM Tris-HCl pH 7.5, 10 mM NaCl, 5 mM MgCl_2_, 1% Triton X-100, 5 mM DTT and protease inhibitor cocktail (protean Mini EDTA-free, Roche)) and lysed using a French press (Thermo Fisher). After ultracentrifugation at 120’000 g for 20 min at 4°C the cleared lysate of GST-tagged GTPases was added to equilibrated glutathione-Sepharose resin (Genescript, USA) and incubated for 1 h at 4°C on a turning wheel. After four washing steps with wash buffer (20 mM Tris-HCl pH 7.5, 10 mM NaCl, 5 mM MgCl_2_) the bound protein was eluted with wash buffer supplemented with 10 mM reduced glutathione (Sigma-Aldrich, Germany).

Cleared lysate of His-tagged GTPases was injected on HisTrap HP columns (GE Healthcare) after equilibration with binding buffer (50 mM Hepes pH 7.5, 150 mM NaCl, 5 mM MgCl_2_, 20 mM imidazole). Washing with 10 column volumes of binding buffer was followed by elution with 5 column volumes of elution buffer (50 mM Hepes pH 7.5, 150 mM NaCl, 5 mM MgCl_2_, 500 mM imidazole). HIS-tagged GTPases were incubated with 50 mM EDTA and further purified by size exclusion chromatography (HiLoad 16/600 Superdex 75 pg, GE Healthcare) with SEC buffer (50 mM Hepes pH 7.5, 150 mM NaCl, 5 mM MgCl_2_, 50 mM EDTA). EDTA was removed by buffer exchange (50 mM Hepes pH 7.5, 150 mM NaCl, 5 mM MgCl_2_) and the protein used for quantitative AMPylation assays.

#### Nucleotide-loading of GTPases

To preload purified GTPases with the respective nucleotide, 50 µM protein was incubated with 3 mM nucleotide (GDP, GTP or GTPγS) and 8 mM EDTA in reaction buffer (50 mM Tris-HCl pH 8.0, 150 mM NaCl, 5 mM MgCl_2_) for 20 min at room temperature. To stop nucleotide exchange 16 mM MgCl_2_ (final) was added. The protein was then used for both *in vitro* AMPylation assays.

#### Radioactive AMPylation assay

*In vitro* AMPylation activity was assayed using either cleared bacterial lysates expressing full-length Bep1 or purified FIC domains of Bep1, VopS and IbpA.

To analyse the AMPylation activity of Bep1, Bep1_FIC_*, VopS_FIC_ and IbpA_FIC2_, 10 µM purified GTPase, preloaded with respective nucleotide, was incubated in presence of the respective AMPylator with 10 µCi [α-^32^P]-ATP (Hartmann Analytic) in reaction buffer (50 mM Tris-HCl pH 8.0, 150 mM NaCl, 5 mM MgCl_2_ containing 0.2 mg/ml RNaseA) for 1 h at 30°C. The reaction was stopped by addition of SDS-sample buffer and heating to 95°C for 5 min. Samples were separated by SDS-PAGE, and subjected to autoradiography.

For AMPylation of Rac1, Cdc42 and their mutant variants, 5 µM of purified HIS-tagged GTPases, preloaded with GDP, were incubation with Bep1_FIC_* (1 µM and 5 µM in Rac1 and Cdc42 variants, respectively) in the presence of [α-^32^P]-ATP for 40 min in reaction buffer (50 mM Tris-HCl pH 8.0, 150 mM NaCl, 5 mM MgCl_2_) at 20°C.

#### Quantitative AMPylation assay

We developed online ion exchange chromatography (oIEC) assay, monitoring the UV absorption of GTPase targets at 260 nm. The observed increase in absorbance due to AMPylation could be readily quantified and resulted in progress curves that yielded reaction velocities and in turn AMPylation efficiencies (*k*_*cat*_*/K*_*M*_).

A 1 ml Resource Q column (GE Healthcare) was equilibrated with loading buffer (20 mM Tris/HCl pH 8.5 or 6.5 for Rac1 or Cdc42, respectively). The purified GTPase variant was mixed with Bep1_FIC_* in reaction buffer (50 mM Tris-HCl pH 7.5, 150 mM NaCl, 5 mM MgCl_2_) in a large volume (200 µl) and the reaction was started at *t* = 0 by addition of 3.2 mM ATP (final concentration, supplemented with 6.4 mM MgCl_2_). A small fraction (20 µL) of the reaction mixture was injected automatically on the column at intervals of 6 minutes. After washing with loading buffer, a gradient of elution buffer (1 M (NH4)_2_SO_4_ in loading buffer) was applied, yielding a chromatogram for each injection. Reaction progress was monitored by quantification of GTPase peak area measured at 260 nm from each chromatogram by numerical peak integration. Note that this peak comprised both native and AMPylated GTPase. A heuristic quadratic function was fitted to the progress curves to yield the initial velocity. Calibration with ATP samples of known concentrations allowed to derive absolute AMPylation velocities. Enzymatic K_M_ and k_cat_ parameters were derived from v_init_(S) type Michaelis-Menten plots (see Fig. S4F and G). Depending on the activity, Bep1_FIC_* concentrations were chosen such that the enzyme velocities were kept within a similar range (see Fig S4H and I). Nominal GTPase concentrations were corrected based on the back-extrapolated peak absorbance at *t* = 0. Fitting of single-substrate kinetic measurements by the Michaelis-Menten equation was developed in python3 with standard modules provided in the Anaconda distribution.

#### Crystallization and structure determination

The full-length biaA gene that codes for the small ORF directly upstream of bep1 gene and part of the bep1 gene from *Bartonella rochalimae* encoding the FIC domain (amino acid residues 13-229) were PCR amplified from genomic DNA. The PCR products for biaA and the fragment of bep1 were cloned into the vector pRSF-Duet1. pRSF-Duet1 containing biaA or bep1 were introduced into *E. coli* BL21 (DE3) by transformation. The constructs were expressed and purified as described for VbhA/VbhT(FIC) (Engel, Goepfert et al., 2012) with the difference that 5 mM DTT was additionally used throughout the purification procedure. Fractions were pooled and concentrated to 13.6 mg/ml for crystallization. Crystals were obtained at 4°C using the hanging-drop vapour diffusion method upon mixing 1 µl protein solution with 1 µl reservoir solution. The reservoir solution was composed of 0.2 M HEPES (pH 7.5), 2.3 M ammonium sulphate and 2% v/v PEG 400. For data collection, crystal was frozen in liquid nitrogen without additional cryoprotectant.

Diffraction data were collected on beam-line X06SA (PXIII) of the Swiss Light Source (λ= 1.0 Å) at 100K on a MAR CCD detector. Data were processed with XDS and the structure solved by molecular replacement with Phaser (McCoy et al., 2007) using the VbhA/VbhT(FIC) structure (PDB ID 3shg) as search model. Several rounds of iterative model building and refinement were performed using Coot (Emsley et al., 2010) and Buster (Bricogne et al., 2016), respectively. The final structure shows high similarity to the VbhA/VbhT(FIC) structure (rmsd 1.44 Å for 183 Cα positions). Crystallographic data are given in Table S2. Figures have been generated using Pymol (2015).

#### Homology modelling of the Bep1:target complex and generation of structure based sequence alignments

The input structure for homology modelling was chosen from all available Rac subfamily structures (i.e., Rac1-3 and RhoG). In total, 43 PDB-entries were analysed (Table S3). Cdc42 (chain D) of the IbpA-Cdc42 complex served as reference for all superimpositions. The superimposition was carried out in two steps: A global superimposition over all Cα atom positions and second, local superimposition using all atom positions of the residues 27-37 (Sw1) of Cdc42. Both steps used the align–algorithm implemented in Pymol (version 1.8) with standard settings. We observed high structural agreement between Rac subfamily GTPase structures in the PDB and the reference chain with an average CαRMSD below 0.5Å. In contrast, we noticed large variations in the all-atom RMSDs of residues in the Sw1 region, that correlate with the nucleotide state of the GTPase. In order to find the most suitable PDB for homology modelling we searched for the smallest coordinate deviations to the Sw1 conformation of the Cdc42 reference chain: Three GDP-loaded GTPase structures display a RMSD of coordinates to the template below 1Å (Table S3). Two of these structures are complexes of the Rho-GDP-dissociation inhibitor (RhoGDI) with either Rac1 (PDB: 1hh4) or Rac2 (PDB: 1ds6) representing the cytosolic “storage form” of the GTPases. The third structure is the Zn^2+^-bound trimeric form of Rac1 (PDB: 2P2L), in which Sw1 is involved in the Zn^2+^-mediated trimer interface. From these candidate PDBs, we chose 1ds6 as the most appropriate for homology modelling, since it represents a physiological state of a Rac-GTPase (in contrast to 2P2L). Further, 1ds6 features a fully resolved Sw1-region and a higher resolution compared to entry 1hh4. To correspond closely to the reference structure, we built an alternative standard rotamer for the solvent-exposed Y32 of Rac2 in the PDB 1ds6 (Fig. 3C). The FIC domains of Bep1 and IbpA were superimposed using the Cα atom positions of flap residues that adopt a β-sheet like conformations in order to mimic the catalytically active conformation of the IbpA:Cdc42 complex. Superimposing IbpA_FIC2_ residues 3667-3670 and 3673-3677, corresponding to Bep1 residues 110-113 and 122-126, respectively yields a rms error of 0.87Å for 9 Cα pairs. Modelling of the complex structure was carried out using the manually selected, superimposed and curated model described above as starting structure for an adapted flexDDG protocol (Barlow et al., 2018) implemented in the Rosetta package. In short: Ligands (GDP and hydrated Mg^2+^) and ordered water molecules (as found in PDB entry 1ds6, as well as 1 water molecule in the center of the Bep1 flap, shown in Figure 2B), that are part of the protein complex interface were parameterized for the use in Rosetta and included in the modelling process to increase precision and validity of the resulting models. The selected small molecules had been refined with B-factors that are comparable to neighbouring mainchain atoms in the respective PDB entries (1ds6 and 5eu0). Next, the curated input model is subjected to a global minimization of backbone and side chain torsions in Rosetta (Minimize step) followed by local sampling of backbone and sidechain degrees of freedom for all residues with C-β atoms within 10Å distance of Rac2 residue D124 (Backrub step). The side chains of the resulting models are optimized globally (Packing step) and backbone and side chain torsion energies are minimized globally (Minimize Step 2). Finally, models are scored on the all atom level using the suggested talaris_2014 function (Barlow et al., 2018) and best scoring models were analysed visually. The recommended total of 35 independent simulations is calculated for the complex with a maximum number of 5000 minimization iterations (convergence limit score 1.0) and 35000 backrub trial steps each. Structure guided multiple sequence alignments (MSA) were generated by manual adjustment of MSA generated using the ClustalW algorithm as implemented in the GENEIOUS software package (Kearse et al., 2012) version 7.1.7.

### Quantification and statistical Analysis

Statistical parameters are indicated in figures and respective legends. Error bars in quantitative AMPylation assays show the standard deviation of reaction efficiencies (k_cat_/K_M_) derived from the least-square minimization of the fitting routine.

### Data and software availability

Accession numbers for the most important structures used in this study are as following: Bep1_FIC_:BiaA (PDB ID: 5EU0, *B. rochalimae* Bep1 uniprot ID: E6YJU0 and BiaA uniprot ID: E6YLF5), IbpA_FIC2_:Cdc42 (PDB ID: 4ITR) and Rac2 (PDB ID: 1ds6). Data analysis for quantitative AMPylation assays was performend in-house with python3. Source code will be made available upon reasonable request.

## Supplemental information

**Figure S1.**
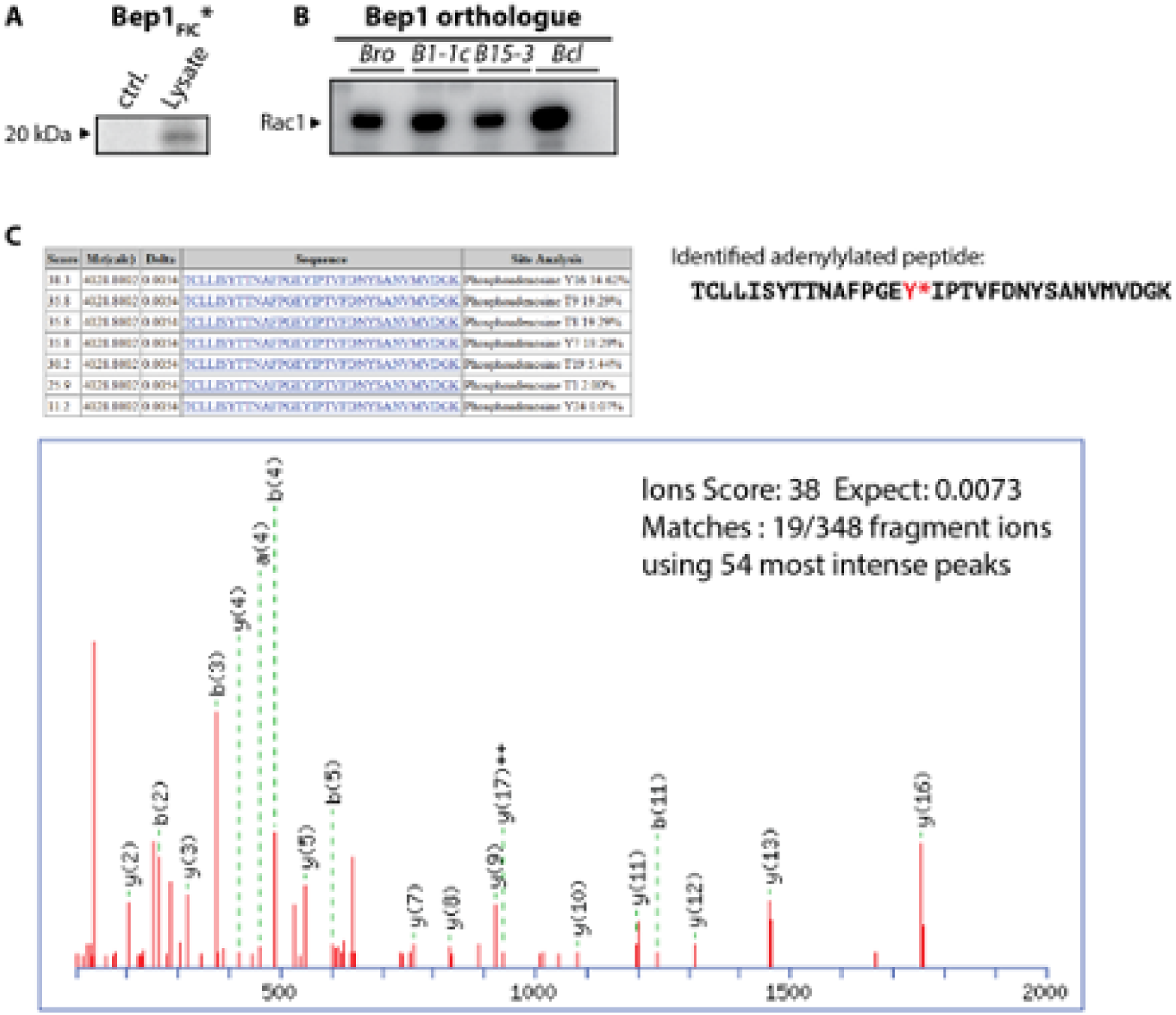
Relates to Figure 1. The Flap of Bep1 is well conserved in orthologues. (**A**) Autoradiograms of Bep1 AMPylation reactions with [α-^32^P] labelled ATP. Bep1 AMPylates an approximately 20 kDa target in J774 cell lysate, indicative of modification of a small GTPase. Lane labelled ctrl. shows no signal for Bep1 only. (**B**) *In vitro* AMPylation activity showing conserved function in Bep1 orthologues of *B. rochalimae* (Bro), *Bartonella sp. 1-1c* (B1-1c), *Bartonella sp. AR15-3* (B15-3), Bartonella *clarridgeiae* (Bcl). (**C**) Identification of the modified peptide by mass spectrometry. Sequence of the identified peptide after tryptic digestion carrying the AMPylation site. The modification is located at tyrosine 16 of the peptide (in red and indicated by an asterisk), corresponding to Y32 of Rac1.

**Figure S2.**
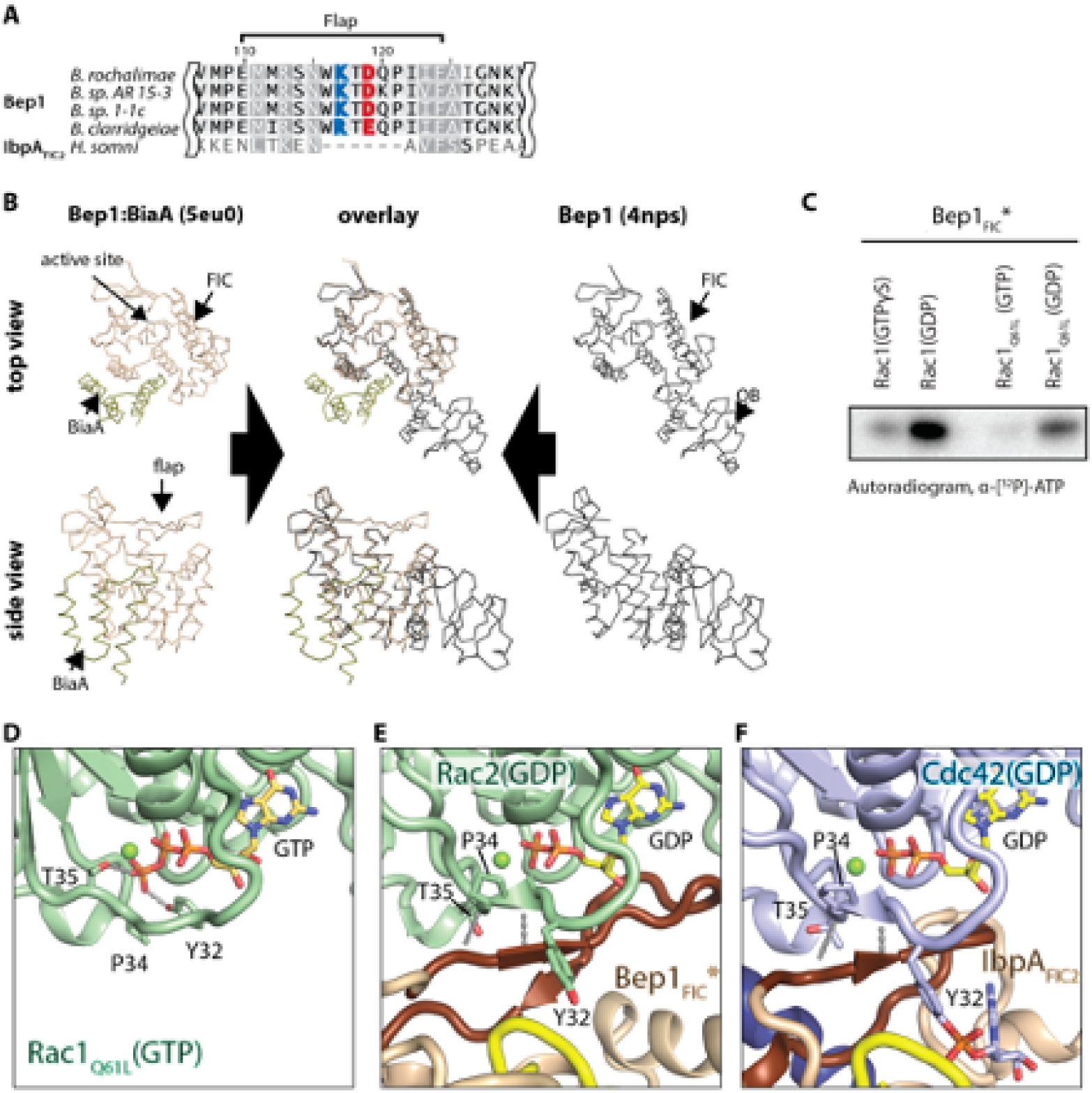
Relates to Figure 2. Bep1_FIC_* is suitable for modelling of a complex with GDP-loaded targets. (**A**) Sequence alignment of flaps found in Bep1 orthologues and IbpA_FIC2_. (**B**) Analysis of BiaA-induced conformational changes in Bep1_FIC_*. Binding of BiaA_E33G_ to Bep1 does not result in detectable conformational changes in the FIC domain as indicated by very small coordinate differences between free (PDB ID: 4nps of *B. clarridgeiae*) and BiaA-bound Bep1 (PDB ID: 5eu0 of *B. rochalimae*): Cα-Coordinate differences for the entire FIC core comprising helices 1 - 5 is 0.37Å (151 CA pairs in residues 42-192) with a smaller deviation the catalytic core (residues 151-191 comprising FIC helices 4-5, RMSD: 0.23Å, 41 Cα pairs). (**C**) Nucleotide dependence of FIC-mediated Rho GTPase AMPylation. Significant Bep1-mediated AMPylation is observed for GDP-loaded Rac1, but not for GTPγS-loaded Rac1 or GTP-loaded, hydrolysis deficient, Rac1_Q61L_ mutant (crystal structure shown in panel (**D**)). Conformation of the switch 1 (Sw1) loop in crystal structures of (**D**) GTP-bound Rac1_Q61L_ and (**E**) GDP-bound Rac2 modelled in complex with Bep1 (GTPase PDB codes are 1e96 and 1ds6, respectively). Notably, Sw1 is in an inward facing conformation in the GTP-bound state shown in (**D**). Y32 is coordinated by the γ-phosphate of the GTPase-bound nucleotide (hydroxyl groups in hydrogen-bonding distance) and is thus inaccessible for modification. In contrast, Sw1 adopts an outward facing conformation in the GDP-bound state shown in (**E**), rendering Y32 solvent accessible. (**F**) Conformation of Sw1 in the product complex between IbpA_FIC2_ and Cdc42 in the GDP-bound state that permits the interaction.

**Figure S3.**
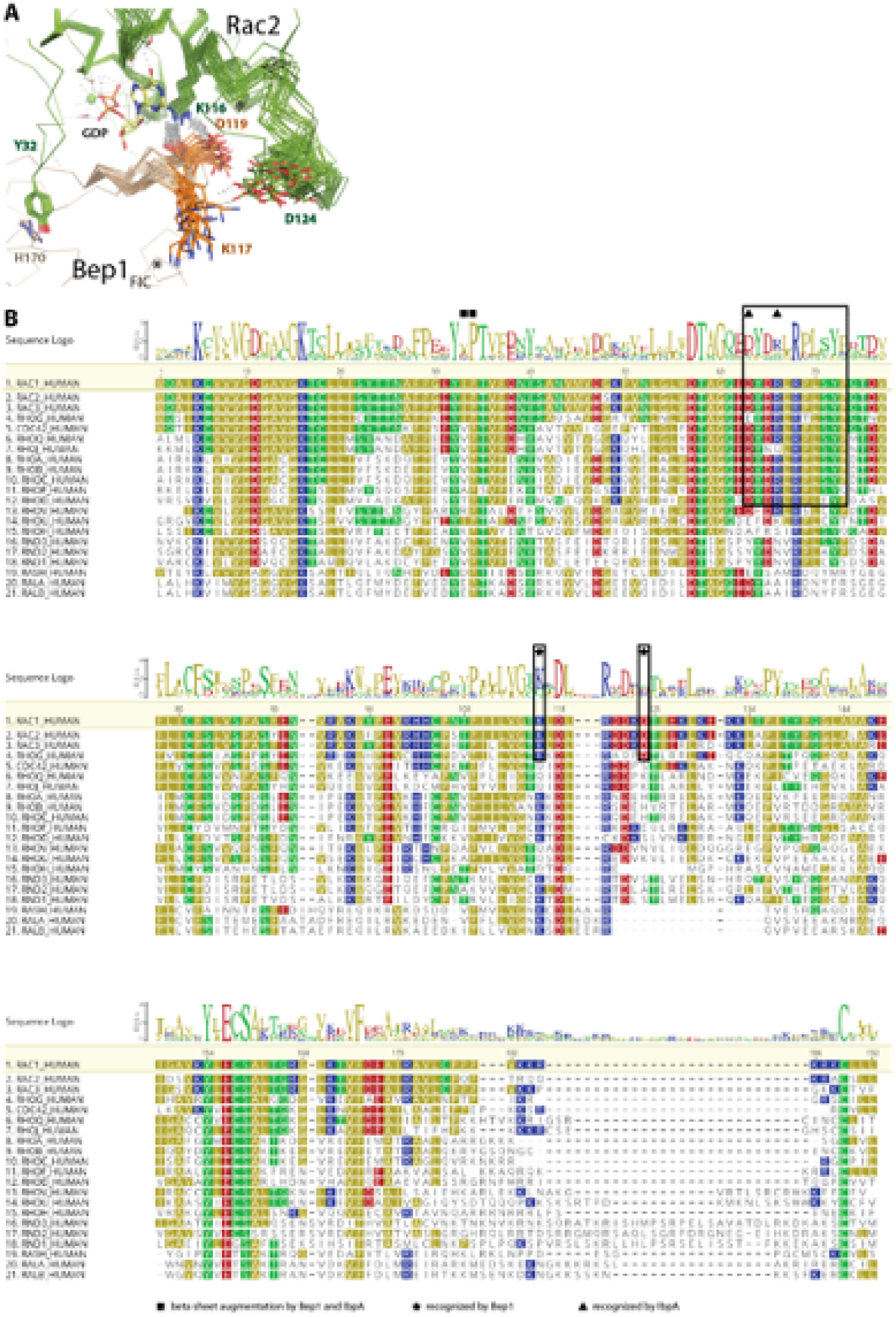
Relates to Figure 3. Proposed target interaction sites for FIC mediated AMPylation. (**A**) Ensemble of Bep1_FIC_:Rac2 models. Bep1_FIC_ (beige) and Rac2 (green) backbones are drawn as wires. Important residues are drawn as sticks. Salt-bridges between Bep1_D119_:Rac1_K116_ and Bep1_K117_:Rac1_D124_ are indicated as dotted lines (grey). Decoys of 25 representative calculations are shown. (**B**) Structure based protein sequence alignment of Rho-GTPases. Side-chain specific interactions with IbpA and Bep1 are indicated by triangles and asterisks, respectively. Interfaces between IbpA and Bep1 and their targets are illustrated as rectangular frames. Residues involved in β-sheet augmentation are marked with squares. Rac1 is set as reference sequence. Polar residues are coloured in green, negatively charged residues in red, positively charged residues in blue and hydrophobic residues in olive.

**Figure S4.**
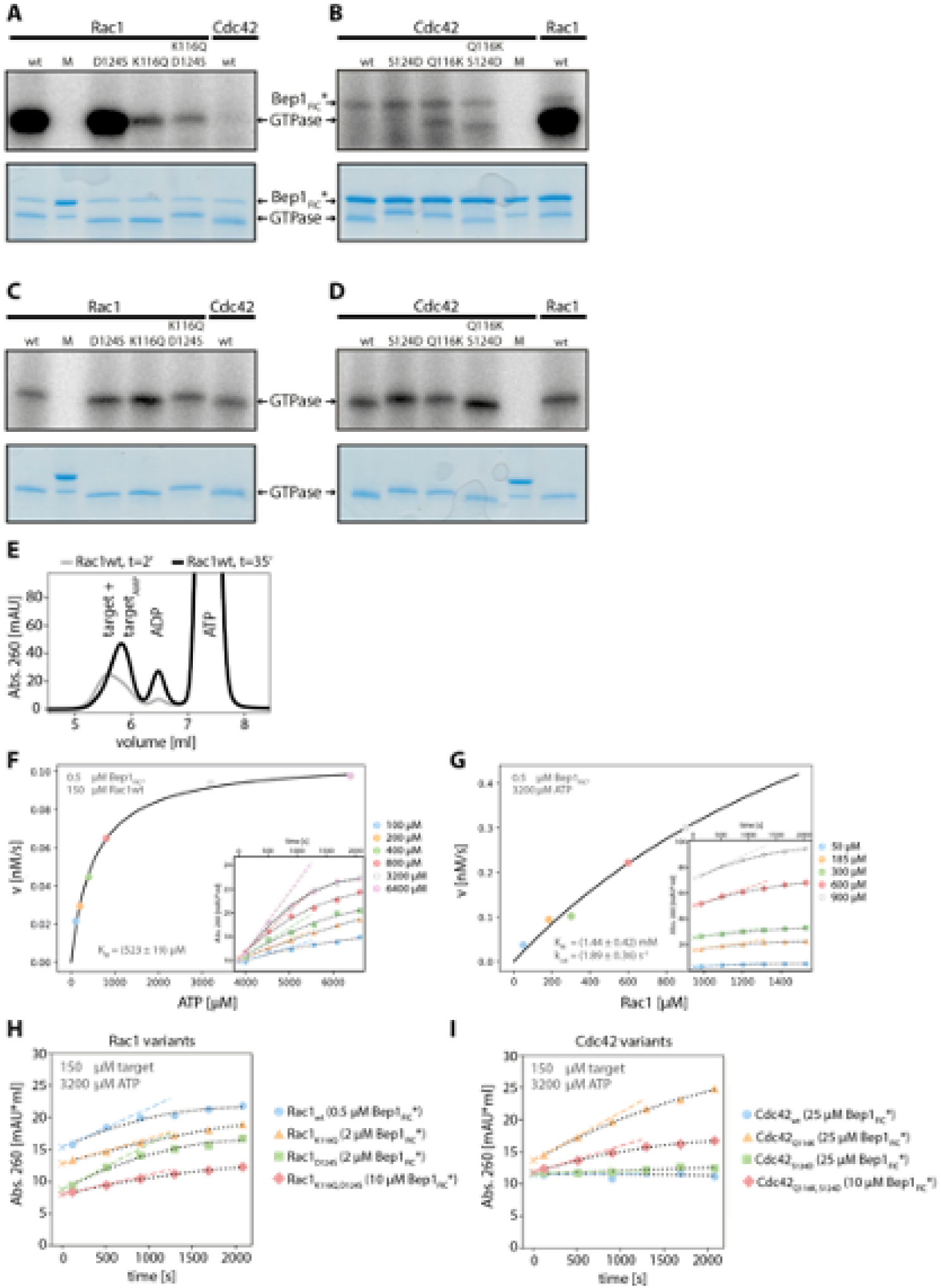
Relates to Figure 4. Raw data showing FIC–mediated AMPylation of GTPase variants. (**A-D**) Autoradiograms and SDS-gels of Rac1 and Cdc42 variants after incubation with [α-^32^P]-labelled ATP and respective AMP-transferases for 40 minutes. (**A**) Rac1 variants and (**B**) Cdc42 variants after incubation with Bep1_FIC_*. (**C**) Rac1 variants and (**D**) Cdc42 variants after concurrent incubation with IbpA_FIC2_. 25 kDa and 20 kDa bands of Precision Plus Protein Standard (Bio-Rad) are visible in all SDS-gels between Rac1_wt_ and the rest of the GTPase variants (lanes labeled ‘M’). (**E**) Ion exchange elution profiles for wild-type Rac1 (Rac1_wt_) at *t* = 2′ (grey) and *t* = 35′ (black), demonstrating the increase in target/target-AMP absorption with time. (**F, G**) Michaelis-Menten plots for the Rac1 ampylation reaction. Initial reaction rates as a function of ATP and Rac1 concentration are shown in panels (**F**) and (**G**), respectively. Initial velocities have been derived from the progress curves shown in the insets. (**H, I**) Progress curves of Bep1_FIC_* mediated AMPylation of Rac1 (**H**) and Cdc42 (**I**) variants. Data points show the absorbance at 260 nm of the target/target-AMP peak during the time course. Heuristic fits are indicated as dotted lines (black). Initial velocities are derived from the first derivatives of the fit-function back-extrapolated to *t* = 0 and drawn as dashed lines in respective.

**Table S1.** Reaction efficiencies of FIC-mediated AMPylation of Rho-GTPase variants.

| *Enzyme* | *Target* | $k_{cat}/K_{M, target}$<br>[s<sup>-1</sup>mM<sup>-1</sup>] | $K_{M, ATP}$<br>[mM] | $K_{M, target}$<br>[mM] | $k_{cat}$<br>[s<sup>-1</sup>] |
| --- | --- | --- | --- | --- | --- |
| **VopS<sub>FIC</sub>** | **Cdc42<sub>Q61L</sub>** | 100 ± 25 <sup>4</sup> | 0.160 ± 0.02 <sup>1</sup> | 0.180 ± 0.04 <sup>1</sup> | 18 ± 1.5 <sup>1</sup> |
| **IbpA<sub>FIC2</sub>** | **Cdc42<sub>Q61L</sub>** | 162 ± 19 <sup>4</sup> | 0.73 ± 0.04 <sup>2</sup> | 1.57 ± 0.15 <sup>2</sup> | 255 ± 15 <sup>2</sup> |
| **Bep1<sub>FIC</sub>\*** | **Rac1<sub>wt</sub>** | 1.31 ± 0.46 <sup>4</sup> | 0.52 ± 0.02 <sup>3</sup> | 1.44 ± 0.42 <sup>3</sup> | 1.89 ± 0.36 <sup>3</sup> |
|  | **Rac1<sub>wt</sub>** | 1.18 ± 0.20 <sup>5</sup> |  |  |  |
|  | **Rac1<sub>D124S</sub>** | 0.481 ± 0.067 <sup>5</sup> |  |  |  |
|  | **Rac1<sub>K116Q</sub>** | 0.202 ± 0.009 <sup>5</sup> |  |  |  |
|  | **Rac1<sub>K116Q, D124S</sub>** | 0.043 ± 0.002 <sup>5</sup> |  |  |  |
|  | **Cdc42<sub>Q116K, D124S</sub>** | 0.046 ± 0.003 <sup>5</sup> |  |  |  |
|  | **Cdc42<sub>Q116K</sub>** | 0.028 ± 0.002 <sup>5</sup> |  |  |  |
|  | **Cdc42<sub>S124D</sub>** | 0.001 ± 0.002 <sup>5</sup> |  |  |  |

<sup>1</sup> taken from (Luong et al., 2010)
<sup>2</sup> taken from (Mattoo et al., 2011)
<sup>3</sup> derived from Figs. S4G and F
<sup>4</sup> derived from $k_{cat}$ and $K_{M, target}$
<sup>5</sup> derived from $v_{init}$ values measured by oIEC (see Figs. S4H and I).

**Table S2.**
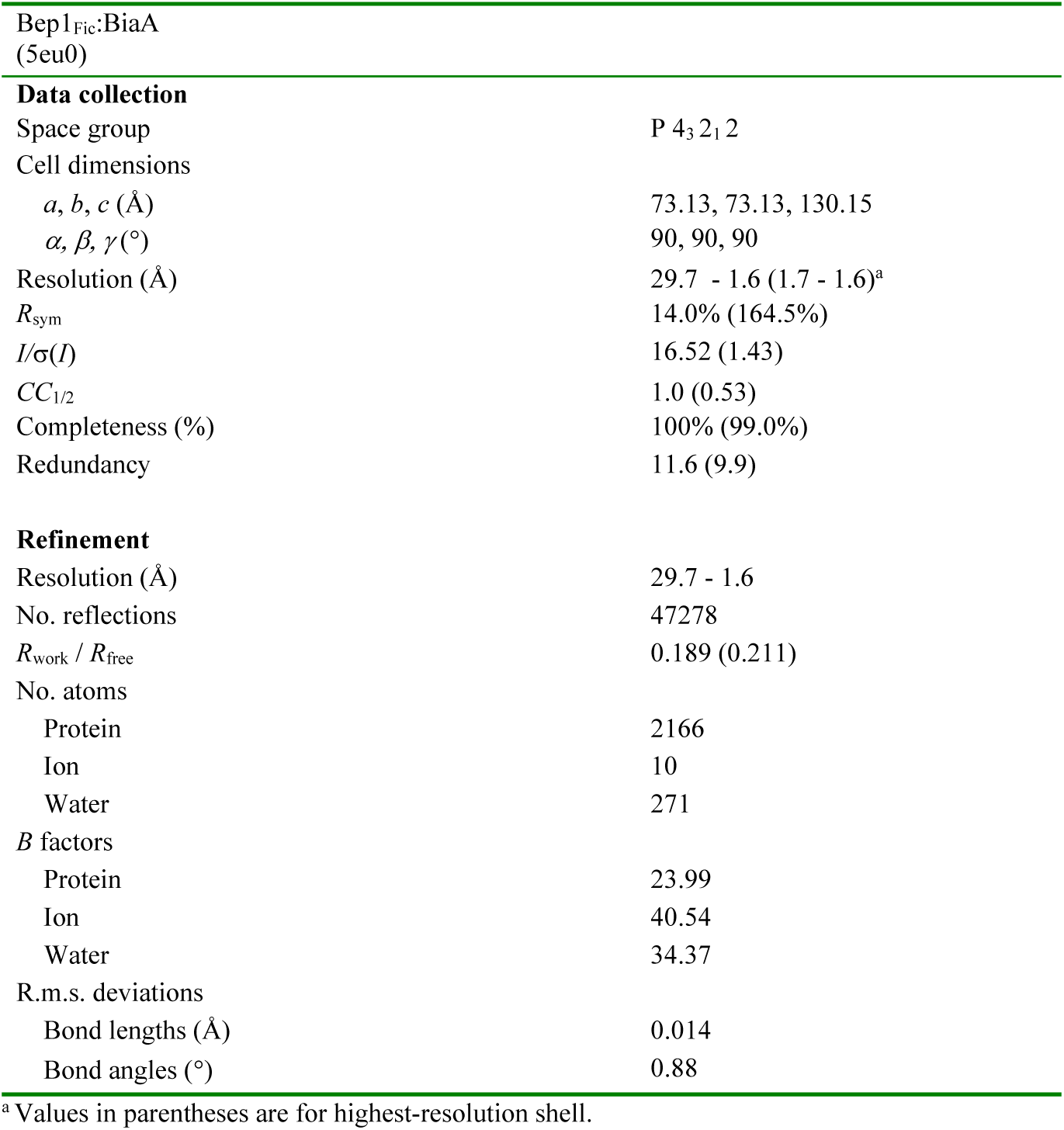
Crystallographic data collection and refinement statistics.

**Table S3.** Global and local structural alignment of Rac-subfamily GTPases to AMPylated Cdc42 in the IbpA bound complex (chain B of PDB entry 4ITR). Chain A of PDB entry 1DS6 was chosen for complex modelling.

| NUCLEOTIDE<br>(ANALOGUE) | PDB ID | GLOBAL<br>RMSD<br>(CA)[Å] | # PAIRS<br>(CA) | USED | LOCAL<br>RMSD<br>(SW1) [Å] | # PAIRS<br>(ATOMS) | USED |
| --- | --- | --- | --- | --- | --- | --- | --- |
| GDP | 1HH4 | 0.534 | 177 | 158 | 0.526 | 85 | 61 |
| GDP | 2P2L | 0.420 | 172 | 133 | 0.781 | 91 | 72 |
| **GDP** | **1DS6** | **0.439** | **175** | **151** | **0.827** | **91** | **65** |
| GDP | 2H7V | 0.453 | 175 | 139 | 1.050 | 91 | 75 |
| GDP | 2W2T | 0.412 | 173 | 144 | 1.867 | 81 | 68 |
| GDP | 2G0N | 0.335 | 175 | 133 | 1.891 | 82 | 67 |
| GDP | 2C2H | 0.411 | 165 | 132 | 1.955 | 48 | 42 |
| GDP | 1I4D | 0.517 | 172 | 135 | 2.419 | 91 | 72 |
| GDP | 1I4L | 0.613 | 173 | 152 | 2.511 | 91 | 77 |
| GDP | 1RYF | 0.392 | 161 | 124 | 3.693 | 89 | 87 |
| GSP | 2W2V | 0.383 | 171 | 138 | 1.973 | 91 | 74 |
| GSP | 2W2X | 0.558 | 167 | 135 | 3.546 | 87 | 77 |
| GSP | 4GZM | 0.492 | 174 | 139 | 4.116 | 91 | 89 |
| GSP | 2FJU | 0.391 | 173 | 130 | 4.169 | 91 | 87 |
| GNP | 2IC5 | 0.357 | 175 | 133 | 1.608 | 88 | 66 |
| GNP | 1I4T | 0.595 | 173 | 147 | 2.520 | 91 | 74 |
| GNP | 1RYH | 0.405 | 162 | 126 | 2.759 | 79 | 68 |
| GNP | 1MH1 | 0.576 | 174 | 144 | 3.191 | 81 | 72 |
| GNP | 3SU8 | 0.633 | 176 | 152 | 3.449 | 91 | 79 |
| GNP | 3RYT | 0.652 | 173 | 154 | 3.488 | 91 | 80 |
| GNP | 3SUA | 0.607 | 176 | 148 | 3.505 | 91 | 79 |
| GNP | 3SBD | 0.416 | 172 | 134 | 3.531 | 91 | 82 |
| GNP | 3TH5 | 0.373 | 168 | 134 | 3.960 | 91 | 87 |
| GNP | 4GZL | 0.449 | 168 | 132 | 4.359 | 82 | 82 |
| GTP | 2WKP | 0.428 | 168 | 143 | 3.317 | 91 | 78 |
| GTP | 1E96 | 0.435 | 176 | 136 | 3.408 | 91 | 79 |
| GTP | 5HZH | 0.463 | 130 | 113 | 3.515 | 78 | 71 |
| GTP | 2WKQ | 0.434 | 168 | 138 | 3.522 | 87 | 77 |
| GTP | 3SBD | 0.416 | 172 | 134 | 3.531 | 91 | 82 |
| GTP | 2WKR | 0.438 | 175 | 143 | 3.703 | 91 | 82 |
| GTP | 1G4U | 0.602 | 172 | 146 | 3.770 | 91 | 84 |
| GTP | 1HE1 | 0.486 | 172 | 142 | 3.912 | 91 | 86 |
| GTP | 4GZM | 0.492 | 174 | 139 | 4.116 | 91 | 89 |
| GTP | 4GZL | 0.449 | 168 | 132 | 4.359 | 82 | 82 |
| GCP | 2QME | 0.517 | 175 | 138 | 3.802 | 87 | 80 |
| GCP | 2OV2 | 0.466 | 172 | 129 | 4.316 | 91 | 91 |
| APO | 2NZ8 | 0.397 | 168 | 122 | 1.625 | 91 | 78 |
| APO | 1FOE | 0.483 | 169 | 131 | 1.651 | 91 | 79 |
| APO | 2VRW | 0.443 | 169 | 133 | 1.691 | 91 | 78 |
| APO | 5FI0 | 0.489 | 169 | 141 | 1.694 | 91 | 79 |
| APO | 4YON | 0.388 | 167 | 125 | 1.765 | 91 | 82 |
| APO | 3BJI | 0.682 | 168 | 138 | 1.871 | 85 | 80 |
| APO | 2YIN | 0.547 | 161 | 139 | 3.476 | 91 | 73 |
| APO | 3B13 | 0.472 | 161 | 137 | 3.645 | 91 | 75 |

